# COVID-19 Mortality is Associated with Impaired Innate Immunity in Pre-existing Health Conditions

**DOI:** 10.1101/2021.05.31.446476

**Authors:** Matthew E. Lee, Yung Chang, Navid Ahmadinejad, Crista E. Johnson-Agbakwu, Celeste Bailey, Li Liu

## Abstract

**Background:** COVID-19 poses a life-threatening endangerment to individuals with chronic diseases. However, not all comorbidities affect COVID-19 prognosis equally. Some increase the risk of COVID-19 related death by more than six folds while others show little to no impact. To prevent severe outcomes, it is critical that we comprehend pre-existing molecular abnormalities in common health conditions that predispose patients to poor prognoses. In this study, we aim to discover some of these molecular risk factors by associating gene expression dysregulations in common health conditions with COVID-19 mortality rates in different cohorts.

**Methods:** We focused on fourteen pre-existing health conditions, for which age-and-sex-adjusted hazard ratios of COVID-19 mortality have been documented. For each health condition, we analyzed existing transcriptomics data to identify differentially expressed genes (DEGs) between affected individuals and unaffected individuals. We then tested if fold changes of any DEG in these pre-existing conditions were correlated with hazard ratios of COVID-19 mortality to discover molecular risk factors. We performed gene set enrichment analysis to identify functional groups overrepresented in these risk factor genes and examined their relationships with the COVID-19 disease pathway.

**Results:** We found that upregulated expression of 70 genes and downregulated expression of 181 genes in pre-existing health conditions were correlated with increased risk of COVID-19 related death. These genes were significantly enriched with endoplasmic reticulum (ER) function, proinflammatory reaction, and interferon production that participate in viral transcription and immune responses to viral infections.

**Conclusions:** Impaired innate immunity in pre-existing health conditions are associated with increased hazard of COVID-19 mortality. The discovered molecular risk factors are potential prognostic biomarkers and targets for therapeutic interventions.

## Background

COVID-19, under devastating inscrutability, was declared a global pandemic by the World Health Organization (WHO) as of March 11, 2020 [1]. The outbreak has seen over 127 million cases and over 2.7 million deaths, and these numbers are still rising [2]. The clinical spectrum of illness ranges from asymptomatic or mild infection to severe pneumonia and death. Well documented risk factors for COVID-19 severity and fatality include age, sex, race, social determinants, and pre-existing health conditions [3–9]. The fatality rate stratified by age groups steadily increases from 0.2 per 100,000 patients in children (<14 years old) to 1797.8 in elders (≥85 years) [4]. Age-adjusted fatality rate in men is 1.4 times higher than in females [5]. More importantly, people with pre-existing health conditions are susceptible to extreme outcomes. While diagnosis rates of COVID-19 are a nearly equal split between patients with and without comorbidities, those with a comorbidity account for 83.29% of COVID-19 deaths [6]. Not all comorbidities have the same impact on COVID-19 prognosis. By linking primary care records of >17 million adults to 10,926 COVID-19 related deaths, the OpenSAFELY project estimated age-and-sex-adjusted hazard ratios (HR) of COVID-19 related deaths for 23 groups of pre-existing health conditions [7]. It reported that patients with organ transplant were at the highest risk (HR=6.00) and those with high blood pressure were at the lowest risk (HR=1.09). However, the biological mechanisms underlying such distinct impacts are largely unknown, which impedes the development of effective interventions to improve clinical outcomes.

Independently, mechanistic studies of COVID-19 pathogenesis have found that dysregulated biological processes in hosts play an important role in disease severity. It is widely recognized that COVID-19 induces cytokine storms with high mortality [10]. Serum levels of inflammatory factors such as interleukins and C-reactive protein have been proposed as prognostic markers [11–13]. Early metabolic responses to infections also show signature differences between patients with favorable outcomes and patients with unfavorable outcomes [14, 15]. Given that comorbidities and molecular dysregulations both influence COVID-19 severity, we must question what molecular abnormalities associated with pre-existing conditions predisposes COVID-19 patients to poor prognosis, and to what extent.

To answer this question, an intuitive approach is to examine molecular profiles of COVID-19 patients before SARS-CoV-2 infection and correlate with prognoses after infection. However, this strategy is infeasible because before-infection samples of COVID-19 patients are rarely collected.

To circumvent this obstacle, we propose to link pre-existing molecular changes to COVID-19 prognosis at the health condition level instead of the individual level, which makes use of summary statistics from epidemiology studies and eliminates the need for individual specific data.

Our strategy is based on the fact that the COVID-19 mortality rate varies with pre-existing conditions. We hypothesize that such variations are associated with molecular changes frequently observed in multiple health conditions. To test this hypothesis, we need quantitative data of COVID-19 mortality rates and molecular dysregulations for various health conditions, which fortunately are readily available. Specifically, the OpenSAFELY study has published the HRs of COVID-19 mortality for 23 groups of pre-existing conditions [7]. For each of these conditions, molecular profiles of affected and unaffected individuals can be found in public repositories, such as the Gene Expression Omnibus (GEO) database [16]. It is widely acknowledged that patient gene expression profiles reflect the underlying pathological changes [17]. While different diseases target different tissues and organs, transcriptomes of peripheral blood cells are informative about systematic changes of a person’s overall health [18, 19]. Therefore, we chose to examine peripheral blood transcriptomes in this study. By correlating transcriptional dysregulations with HRs of COVID-19 mortality, we discovered molecular risk factors that predispose COVID-19 patients to severe outcomes. We further analyzed functional relationships of these risk genes, which revealed known and novel biological mechanisms of COVID-19 pathogenesis.

While we focused on gene expression risk factors in this study, our analytic approach is applicable to other omics-level profiles, such as epigenetic and metabolomic data. Integration of these discoveries will allow for better prediction of severe outcomes of COVID-19 and inform the development of preventative measures to reduce fatality. Furthermore, the long-term sequelae of COVID-19 survivors are currently unknown. A greater apprehension of the disease mechanisms in the context of comorbidities will serve for future evaluation of the health impact of COVID-19 on patients with chronic diseases.

## Methods and Materials

### Data sets

The OpenSAFELY project reported age-sex-adjusted HRs of COVID-19 related deaths for 23 groups of pre-existing health conditions. We excluded the two cancer groups (solid tumors and hematological malignancies) due to the extremely high heterogeneity of cancers [20]. For each remaining health condition, we searched the GEO database [16] to identify transcriptomics studies involving affected individuals (cases) and unaffected individuals (controls, **Fig. 1A**). We limited our queries to peripheral blood samples as a modus operandi of removing confounders related to different tissue types and encapsulating disease characteristics at a systematic level. We further limited our query to microarray-based transcriptomic profiles to reduce technical variance. If multiple data sets were available for a health condition, we chose the one with the largest sample size. We downloaded the normalized gene expression values.

**Figure 1.**
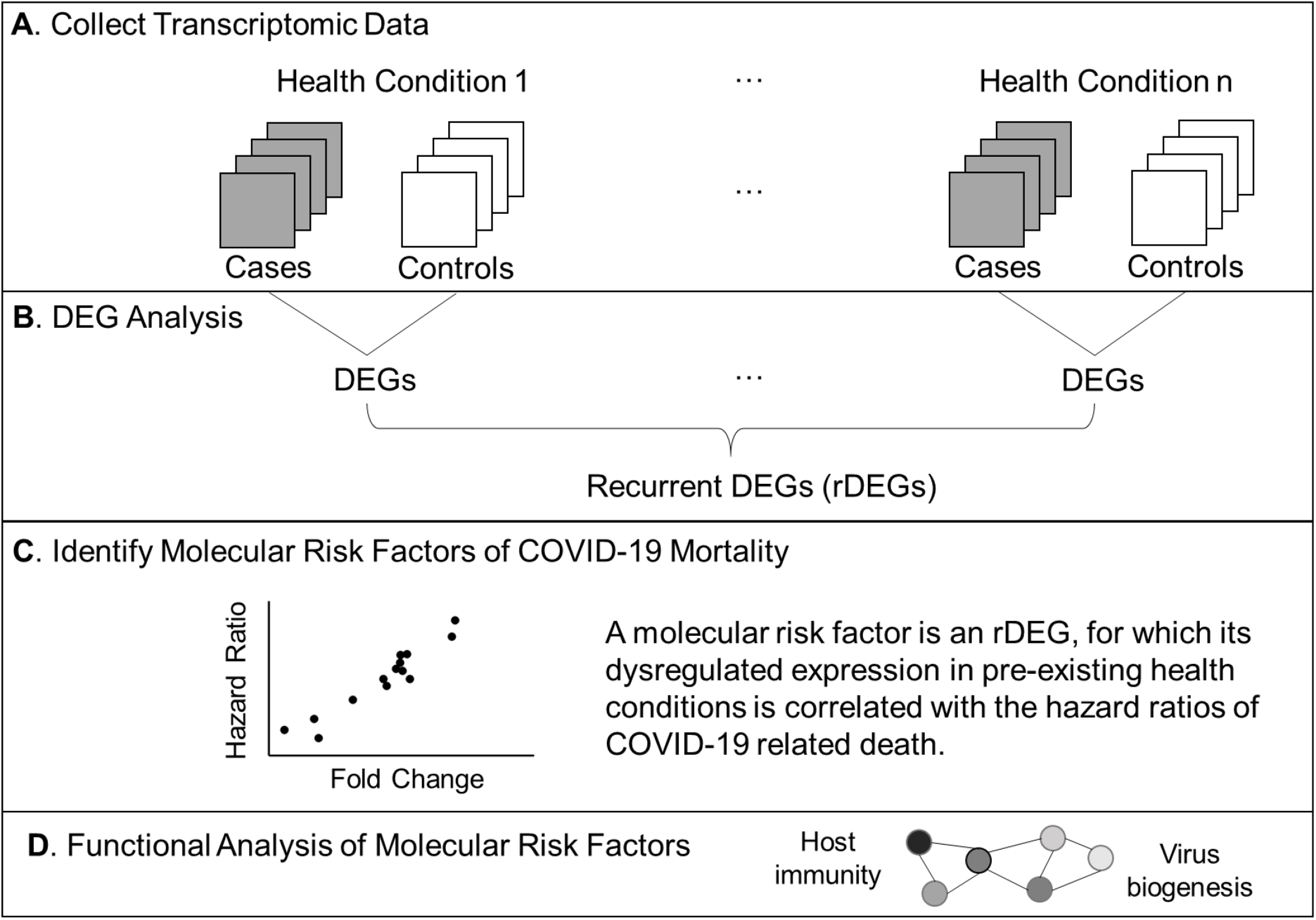
Analysis workflow. (**A**) Compile microarray-based transcriptomic data sets for common health conditions. We included case-control studies using peripheral blood samples. (**B**) Find differentially expressed genes (DEGs) for each health condition. Recurrent DEGs were differentially expressed in at least four health conditions. (**C**) Perform correlation tests to identify molecular risk factors, which are pre-existing expression dysregulations that increase the risk of COVID-19 related death. (**D**) Examine functional relationships of molecular risk factors via enrichment and network analyses.

### Identify dysregulated gene expression in pre-existing health conditions

Given a transcriptomic data set, we used the Student t-test to compare expression levels of each gene in cases versus controls (**Fig. 1B**). Aiming to be inclusive at this progressive state of analysis, we considered genes with nominal P value <0.05 to be differentially expressed. If multiple probes on the microarray represented the same gene, we kept the one with the best P value and removed the other ones to avoid redundancy. For a differentially expressed gene (DEG), we computed the fold change (FC) as the ratio of the mean expression level in cases over controls. For a non-DEG, we set the FC to one. If a gene was differentially expressed in at least four health conditions, it was a recurrent DEG (rDEG).

### Identify molecular risk factors of COVID-19 mortality

For each rDEG, we tested if its FCs in pre-existing conditions were correlated with HRs of COVID- 19 mortality using Pearson correlation test (**Fig. 1C**). We corrected for multiple comparisons by converting nominal P values to false discovery rates (FDRs) using the Benjamini-Hochberg method [21]. FDR<0.05 indicated significant molecular risk factors. FDR>0.05 but nominal P value <0.01 indicated marginal risk factors. The positive sign or the negative sign of Pearson correlation coefficient (PCC) indicated that upregulated gene expression or downregulated gene expression in pre-existing conditions increased the risk of COVID-19 mortality, respectively.

### Functional categorization and analysis

We classified molecular risk factors into overlapping gene sets based on annotations of biological processes in the Gene Ontology database and pathways in the KEGG database. For each gene set, we tested if it was overrepresented using Fisher’s exact test. We corrected for multiple comparisons by converting nominal P values to FDRs. We built association networks of enriched gene sets (FDR<0.05) and examined their relationships (**Fig. 1D**). We used the clusterProfiler and enrichplot packages in R/Bioconductor for these analyses [22].

## Results

### DEGs in pre-existing health conditions

Our query of the GEO database found qualified transcriptomic data for fourteen health conditions. For each health condition, we identified DEGs with Student t-test P <0.05. This lenient cutoff allowed us to be as inclusive as possible at this step. On average, each health condition was associated with 5,777 DEGs (range 1215 to 17,688). Most of the DEGs were downregulated in cases as compared to controls (mean FCs range from 0.003 to 0.160). Among a total of 25,552 genes analyzed, we found 11,930 rDEGs that were differentially expressed in at least four health conditions. **Table 1** presents the summary statistics of DEGs.

**Table 1:** Data sets and DEGs in fourteen pre-existing conditions.

| Health Condition | Gene Expression |  |  | COVID-19 Mortality * |  |
| --- | --- | --- | --- | --- | --- |
|  | GEO Accession | DEG Count | Mean FC | Disease Group | HR |
| Alzheimer's disease | GSE63060 | 5,413 | 0.99 | Stroke or dementia | 2.57 |
| Asthma | GSE110551 | 1,215 | 1.01 | Asthma | 1.55 |
| Coronary artery disease | GSE12288 | 1,289 | 1.10 | Chronic heart disease | 1.57 |
| Chronic obstructive pulmonary disease | GSE42057 | 2,054 | 0.99 | Respiratory disease excluding asthma | 1.95 |
| Hispanic ethnicity | GSE30101 | 4,249 | 1.13 | Ethnicity | 1.37 |
| HIV | GSE104640 | 1,1503 | 1.00 | Immunosuppressive condition | 2.75 |
| Hypertension | GSE135111 | 413 | 1.53 | High blood pressure or diagnosed hypertension | 1.09 |
| Lupus | GSE37356 | 2,289 | 1.00 | Rheumatoid arthritis, lupus or psoriasis | 1.30 |
| Obesity | GSE110551 | 3,905 | 1.00 | BMI >35 | 1.81 |
| Rheumatoid Arthritis | GSE93272 | 9,902 | 1.01 | Rheumatoid arthritis, lupus or psoriasis | 1.30 |
| Type-2 Diabetes | GSE65561 | 4,155 | 0.99 | Diabetes | 2.27 |
| Chronic Kidney Disease | GSE37171 | 17,688 | 1.01 | Reduced kidney function (eGFR<30) | 3.48 |
| Multiple sclerosis | GSE21942 | 7,735 | 1.12 | Other neurological disease | 3.08 |
| Alcoholic Hepatitis | GSE28619 | 9,489 | 1.05 | Liver disease | 2.39 |

\* as reported in the OpenSAFELY Project [7]

### Pre-existing expression dysregulations increase COVID-19 death risks

For each rDEG, we tested if its FCs in different health conditions were correlated with HRs of COVID-19 mortality. For health conditions where this gene was not differentially expressed, we set the FCs to 1 and included them in the correlation test as well. Among a total of 11,930 DEGs, we found no significant molecular risk factor that passed the stringent FDR<0.05 threshold. However, 251 genes passed the Pearson correlation test P <0.01 threshold and were considered as marginal molecular risk factors. Among them, upregulated expression of 70 genes and downregulated expression of 181 genes increased risk of COVID-19 related death (**Supplementary Table 1**).

The *RPS28* gene had the most significant correlation P value (0.0003). Its FCs in pre-existing conditions were positively correlated with HRs of COVID-19 mortality (PCC= 0.83, **Fig. 2A**). *RPS28* encodes a component of the 40S subunit of ribosome where a cell synthesizes proteins. It was differentially expressed in six health conditions, including rheumatoid arthritis, chronic obstructive pulmonary disease, alcoholic hepatitis, multiple sclerosis, HIV, and chronic kidney disease. As its FC increased from 0.98 to 1.14, the HR of COVID-19 mortality increased from 1.30 to 3.48. Furthermore, the list of molecular risk factors contained seven additional genes that encode ribosomal components *(RPLP1, RPLP2, RPL13, RPL23A, RPL30, RPL38,* and *RPS11*). Except *RPL30,* upregulation of these genes consistently increased the HR of COVID-19 mortality (P range= 0.001 to 0.009, PCC range= 0.71 to 0.78, **Fig. 2B**). This is in accordance with the positive viral infection-regulating roles of ribosomal proteins [23]. Notably, three ribosomal proteins in our list are required for early virus accumulation [24], though mechanistic studies in SARS-CoV-2 are still lacking.

**Figure 2:**
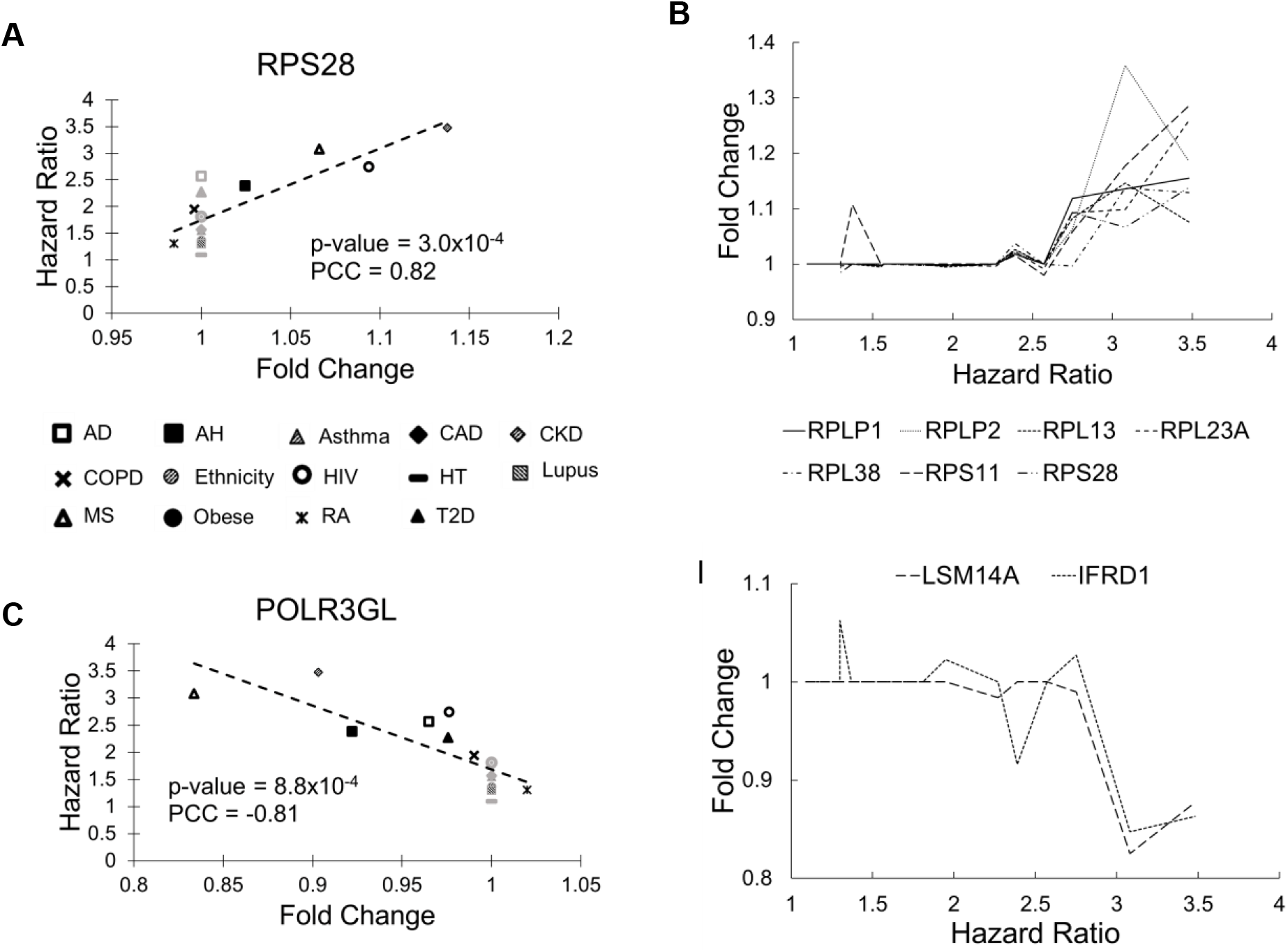
Scatter plots of selected genes showing correlations between FC and HR. (**A**) *RPS28* had the best Pearson correlation test *P* values and positive PCC values. The fourteen health conditions are represented by different symbols (AD: Alzheimer’s disease, AH: alcoholic hepatitis, Asthma: asthma, CAD: coronary artery disease, CKD: chronic kidney disease, COPD: chronic obstructive pulmonary disease, Ethnicity: Latino vs. Caucasian, HIV: human immunodeficiency virus, HT: hypertension, Lupus: lupus, MS: multiple sclerosis, Obese: obesity, RA: rheumatoid arthritis, and T2D: type-2 diabetes). Black symbols indicate health conditions in which a given gene was differentially expressed. Gray symbols indicate health conditions in which a given gene was not differentially expressed. Broken lines represent linear fits between FC and HR. (**B**) Seven genes coding ribosomal proteins consistently show positive correlations between expression FCs and HRs of COVID-19 mortality. (**C**) *POLR3GL* had the best Pearson correlation test *P* values and negative PCC values. It is involved interferon production providing anti-viral innate immunity. (**D**) Two additional genes involved in interferon signaling both show negative correlations between FC and HR.

The *POLR3GL* gene showed the most significant negative correlation (P= 0.0008, PCC= –0.81, **Fig. 2C**). *POLR3GL* encodes a subunit of RNA polymerase III that catalyzes the transcription of DNA into RNA. It induces production of interferon *(IFN-α/β)* to inhibit virus replications [25, 26]. Consistent with this function, pre-existing downregulation of *POLR3GL* in seven health conditions (Alzheimer’s disease, chronic kidney disease, alcoholic hepatitis, chronic obstructive pulmonary disease, HIV, multiple sclerosis, rheumatoid arthritis, and type-2 diabetes) increased the risk of COVID-19 related death. Furthermore, the list of risk factors contained two additional genes *LSM14A* and *IFRD1* that regulate interferon signaling. For both genes, downregulation increased HRs of COVID-19 mortality (P = 0.003 and 0.005, PCC = −0.74 and −0.71, respectively, **Fig. 2D**). Interestingly, we did not find interferons as molecular risk factors.

### Functional groups enriched with risk factors

We classified the list of marginal molecular risk factors into functional gene sets based on Gene Ontology and KEGG annotations and performed enrichment analysis. At FDR<0.05, these molecular risk factors were significantly enriched in ten biological processes and three pathways (**Table 2**). Most of these gene sets were related to viral transcription, mRNA processing and metabolism, protein synthesis, and endoplasmic reticulum (ER) function.

**Table 2:** Significantly enriched gene sets.

| ID | Description | Enrichment | FDR | Count | Genes |
| --- | --- | --- | --- | --- | --- |
| **KEGG Pathways** |  |  |  |  |  |
| hsa03010 | Ribosome | 4.77 | 0.00026 | 8 | RPL38/RPS11/RPS28/RPLP1/RPLP2/RPL13/RPL30/RPL23A |
| hsa00563 | GPI-anchor biosynthesis | 10.87 | 0.0025 | 3 | PIGP/PIGA/PIGG |
| hsa05171 | Coronavirus disease COVID-19 | 3.25 | 0.0031 | 8 | RPL38/RPS11/RPS28/RPLP1/RPLP2/RPL13/RPL30/RPL23A |
| **GeneOntology Biological Process** |  |  |  |  |  |
| GO:0045047 | protein targeting to ER | 8.55 | 0.00036 | 10 | RPL38/RPS11/SEC62/RPS28/RPLP1/RPLP2/SRP9/RPL13/SGTB/RPL23A |
| GO:0006613 | cotranslational protein targeting to membrane | 8.33 | 0.00087 | 9 | RPL38/RPS11/SEC62/RPS28/RPLP1/RPLP2/SRP9/RPL13/RPL23A |
| GO:0022626 | cytosolic ribosome | 7.34 | 0.0040 | 8 | RPL38/RPS11/RPS28/RPLP1/RPLP2/RPL13/RPL30/RPL23A |
| GO:0006413 | translational initiation | 5.23 | 0.0055 | 10 | HABP4/RPL38/RPS11/RPS28/RPLP1/RPLP2/RPL13/P AIP2/COPS5/RPL23A |
| GO:0019083 | viral transcription | 5.13 | 0.013 | 9 | RPL38/RPS11/RPS28/RPLP1/RPLP2/RPL13/NUP160/POLR2C/RPL23A |
| GO:00116607 | nuclear speck | 3.45 | 0.014 | 13 | PNN/HABP4/CBLL1/HBP1/ACADM/SRSF2/SRSF7/RFXAP/DDX42/SART1/POLDIP3/NRIP1/SAP18 |
| GO:0000184 | nuclear-transcribed mRNA catabolic process, nonsense-mediated decay | 5.89 | 0.031 | 7 | RPL38/RPS11/RPS28/RPLP1/RPLP2/RPL13/RPL23A |
| GO:0008380 | RNA splicing | 3.01 | 0.032 | 14 | PNN/HABP4/DHX8/ZRANB2/HNRNPH3/HNRNPA2B1/ZNF326/SRSF2/SRSF7/DDX42/PPWD1/SART1/POLR2C/SAP18 |
| GO:0044391 | ribosomal subunit | 4.32 | 0.040 | 8 | RPL38/RPS11/RPS28/RPLP1/RPLP2/RPL13/RPL30/RPL23A |
| GO:0031307 | integral component of mitochondrial outer membrane | 15.40 | 0.049 | 3 | SYNJ2BP/TOMM20/BNIP3 |

We then built an association network and examined the functional relationships of the enriched gene sets and the molecular risk factors. We observed three clusters (**Fig. 3**). Each cluster is composed of highly interconnected gene sets. Crosstalk between clusters is indicated with individual genes bridging these clusters.

**Figure 3:**
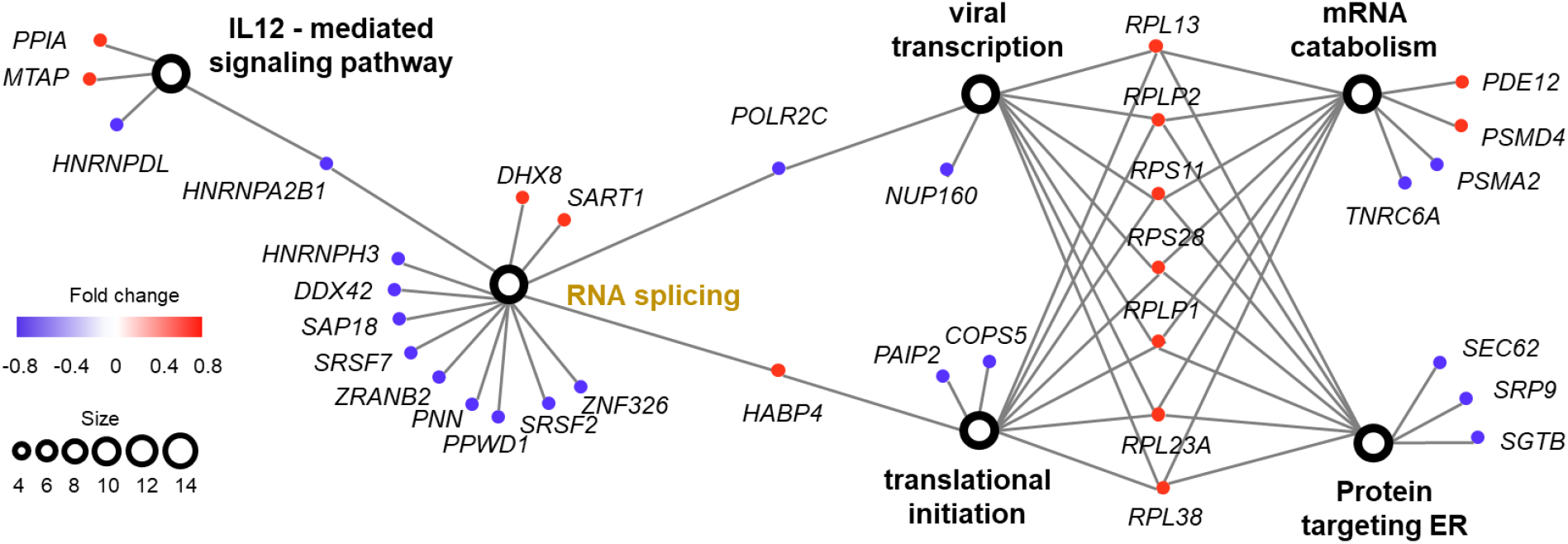
Association networks of functional groups enriched with molecular risk factor genes. This network contained two types of nodes, representing gene sets (open circles) and individual risk factor genes (colored dots). An edge links a risk factor gene to its associated gene set. The size of an open circle is proportional to the number of risk factor genes connected to it. The color of a dot indicates if its upregulation is positively (red) or negatively (blue) correlated with HR of COVID-19 mortality. All six gene sets had FDR<0.05 from enrichment analysis (Table 2).

The first cluster consisted of gene sets involved in viral transcription, translational initiation, mRNA catabolic process, and proteins targeting ER. These gene sets shared eight common risk factor genes encoding ribosomal components that function in ER. Except *RPL30,* upregulation of all these genes increased the HR of COVID-19 mortality. Conversely, for most of the other genes (7 out of 9) in this cluster, downregulation increased the HR of COVID-19 mortality, including two genes *(SEC62* and *SGTB)* that target ER and participate in degradation of misfolded proteins. Therefore, pre-existing abnormal functions in ER, specifically upregulated protein synthesis and downregulated degradation of unfolded protein were associated with high risk of COVID-19 death.

The second cluster consisted of a single gene set involved in RNA splicing. For almost all genes (11 out of 14) in this cluster, downregulation increased the HR of COVID-19 mortality. In COVID- 19 patients, host RNA splicing is significantly disrupted by SARS-CoV-2 [27]. Our observation suggests that pre-existing downregulation of RNA splicing genes can potentially aggravates such disruptions.

The third cluster consisted of a single gene set participating in interleukin-12-mediated signaling pathway. Noticeably, one of the risk factor genes in this cluster, namely *PPIA* has been shown to act as a potential mediator between human SARS coronavirus nucleoprotein and *BSG/CD147* in the process of invasion of host cells by the virus [28]. Consistent with this previous study, we found that pre-existing upregulation of this gene increased COVID-19 mortality risk in nine common health conditions (**Fig. 4A**). Interestingly, interleukin-12 *(IL12A* and *IL12B* genes) was not a molecular risk factor. It was dysregulated in two health conditions but did not meet the criteria of being a rDEG, which required dysregulation in at least four health conditions. Its receptor *IL12RB1* was a near miss, showing positive correlation between FC and HR in five health conditions (P=0.079, PCC=0.48).

**Figure 4:**
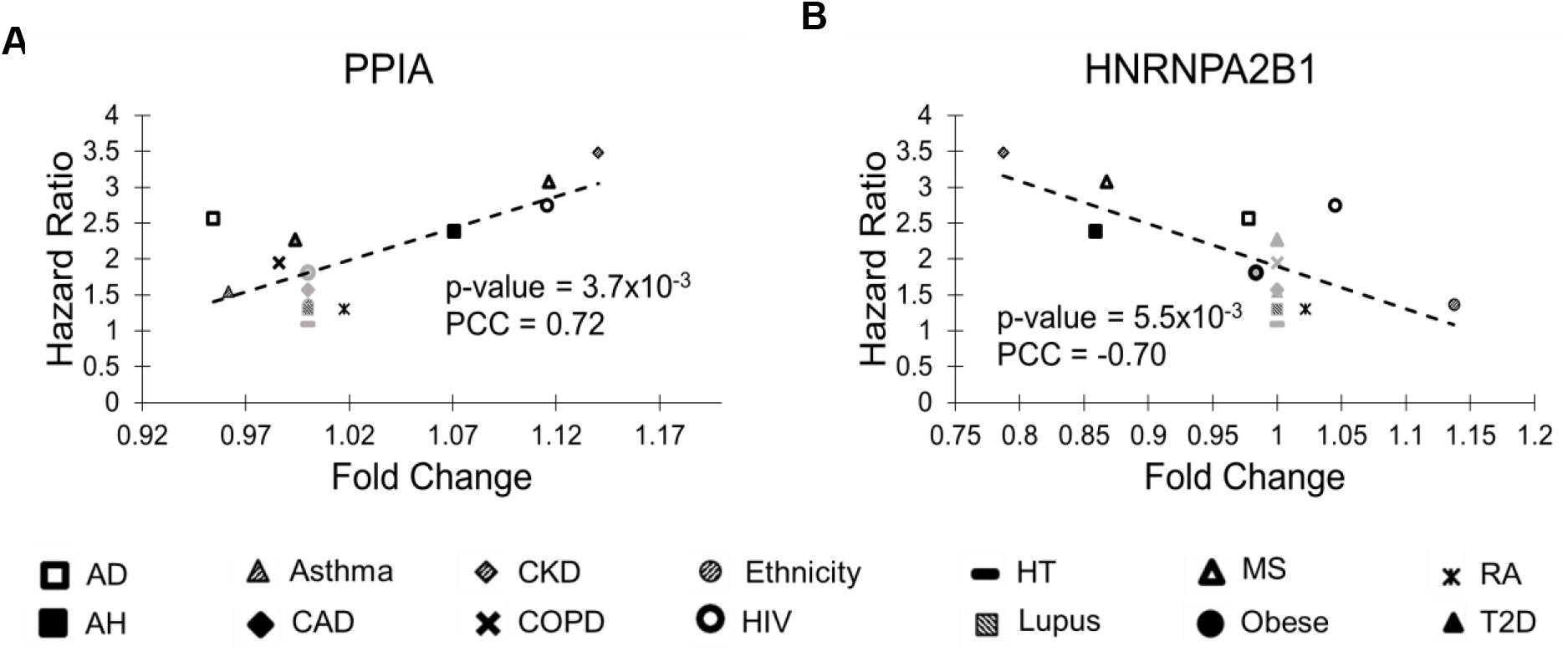
Scatter plots of two genes in the IL12-mediated signaling pathway. (**A**) Upregulation of *PPIA* increased risk of COVID-19 mortality. *PPIA* facilitates virus replication. (**B**) Downregulation of *HNRNPA2B1* increased risk of COVID-19 mortality. *HNRNPA2B1* has anti-virus replication activity. Figure legend is the same as in Fig. 2.

Crosstalk between the second and third clusters was via *HNRNPA2B1* that binds heterogeneous nuclear RNA (hnRNA) and subsequently induces IFN-α/β production to inhibit virus replication. Two other hnRNA binding proteins, *HNRNPH3* and *HNRNPDL,* were also molecular risk factors. For all three genes, pre-existing downregulation increased COVID-19 mortality risk, presumably by blocking IFN-α/β production which compromises the innate immunity (**Fig. 4B**).

## Discussion

The drastically different disease progression and prognosis among COVID-19 patients with preexisting health conditions challenge clinical management of this life-threatening disease. In this study, we integrated publicly available transcriptomics data of common health conditions and COVID-19 epidemiology data to study the molecular mechanisms underlying this complex problem. Our analyses revealed that pre-existing transcriptional dysregulations increased risk of severe COVID-19 outcomes, plausibly via impairing host innate immunity, as discussed below.

The list of molecular risk factors was enriched with genes targeting ER. ER is a crucial organelle controlling various essential physiological functions in eukaryotic cells. Perturbation of ER homeostasis causes ER stress that compromises host immunity and induces cell death via apoptosis [29]. During viral infection, ER is hijacked for entry to host cells and replication and assembly of viral genomes [30]. Emerging evidence suggests that coronavirus infection, including SARS and COVID-19, triggers ER stress that sequentially activates unfolded protein response and inflammatory reactions [31–33]. Indeed, besides the *SEC62* and *SGTB* genes that degrade unfolded proteins in ER, the list of molecular risk factors contains four additional genes *(UBE2J2, COPS5, MBTPS2,* and *PPIA)* that respond to unfolded proteins. Therefore, abnormal ER stress responses in pre-existing health conditions plausibly leads to severe COVID-19 outcomes by exacerbating ER stress during SARS-CoV-2 infection, provoking proinflammatory reactions and accelerating host cell death. This new finding implies potential applications of preventive interventions to reduce ER stress and subsequently improve COVID-19 prognosis. From this perspective, compounds that have been studied to treat ER stress in other diseases are promising candidates [34–36].

Our results also imply that interferon production and signaling are compromised in several health conditions via downregulation of *POLR3GL, LSM14A, IFRD1,* and three hnRNA binding proteins *(HNRNPA2B1, HNRNPH3,* and *HNRNPDL).* Reduced antiviral interferon response has been associated with excessive proinflammatory responses in COVID-19, which leads to severe outcomes [37]. Our analyses thus identified transcriptional dysregulations that predispose patients to poor prognosis by disrupting type I interferon signaling pathways. Meanwhile, IL12- mediated signal pathway is enriched with molecular risk factors. Similar to interferons, IL12 are proinflammatory cytokines that mediate the innate immune response [38]. However, we did not find any cytokines, either proinflammatory or anti-inflammatory, as molecular risk factors. Therefore, our results do not directly explain the association of cytokine storm with COVID-19 severity.

Limitations of our study include the lack of individual risk factors passing a stringent statistical threshold and no consideration of multivariate effects. Although our analysis identified marginal molecular risk factors passing the nominal *P* value cutoff, none had a significant FDR after correction for multiple comparisons, which disqualified them as prognostic markers. However, analysis using the aggregation of these risk factor genes discovered significantly enriched biological processes, with the best FDR<10^-4^ (**Table 2**). Therefore, we are confident that chronic ER stress and immune dysregulation in pre-existing health conditions increased risk of COVID- 19 mortality. Our analyses were based on univariate models, in which we examined the expression levels of each gene separately. Because multiple genes are dysregulated concurrently and a combination of them contributes to COVID-19 prognosis, a more realistic model should consider their combined effect. However, because the transcriptomics data were derived from individual patients and HRs of COVID-19 mortality were from summary statistics of an epidemiology study, we chose to use univariate models that are more straightforward to interpret.

Our novel analytical approach integrates epidemiology data and omics data to discover molecular risk factors. While we focus on transcriptional regulation in this study, an immediate next step is to apply this approach to other molecular changes, including genetic variation, epigenetic modification, and metabolic perturbation to investigate their roles in COVID-19 pathogenesis. As before-infection samples of COVID-19 patients are difficult to acquire, integration of existing multi-omics data and epidemiology data hold the promise to accelerate the discovery of diagnostic and therapeutic markers to improve the management of COVID-19 disease.

## Conclusions

By the harmonic techniques conveyed before, our study illuminates that gene expression dysregulation in pre-existing health conditions that impair innate immunity are molecular risk factors of COVID-19 related death. The individual risk factor genes or gene sets are potential mediators in disease pathogenesis. These findings allow for better prediction of severe outcome, inform the development of preventative measures to reduce fatality, and inform the evaluation of long-term health impact of COVID-19 in different populations.

